# Differences between males and females in psychostimulant/sucrose self-administration (when observed) may not necessarily be driven by biological sex

**DOI:** 10.64898/2026.06.09.731125

**Authors:** Indu Mithra Madhuranthakam, Shakil Ahmed, Kona Basak, Azim Uddin, Mst Afroza Alam Tumpa, Alida Jimenez, Rachel Cherry, Adriana Rodriguez, Maria Chowdhury, Thomas M. Keck, Martin O. Job

**Affiliations:** Department of Chemistry and Biochemistry, Rowan University, 201 Mullica Hill Road, Glassboro, New Jersey, USA; Department of Biomedical Sciences, Cooper Medical School of Rowan University, 401 S Broadway, Camden, New Jersey, USA

**Author notes:** Corresponding author: (MOJ).

**Keywords:** Biological sex, sex differences, psychostimulant, methamphetamine, self-administration, individual differences, sucrose self-administration

## Abstract

**Background:** Sex differences in psychostimulant-related behaviors are often attributed to biological sex; however, individual variability may also strongly influence behavioral outcomes. The new MISSING (Mapping Intrinsic Sex Similarities as an Integral quality of Normalized Groups) model identifies mixed-sex behavioral groups in which differences are driven primarily by individual variability rather than sex. The goal of this study was to validate the MISSING model for psychostimulant/sucrose self-administration.

**Methods:** Long Evans rats self-administered methamphetamine (METH, male n = 25, female n = 32, 0.1 mg/kg/infusion, FR1, 6h per day for 20 days), sucrose (male n = 20, female n = 22, one-20 mg pellet/delivery, all other conditions being equal) and saline (male n = 3, female n = 10, other things being equal). We developed a new Quantitative Structure of Curve Analytical (QSCAn) model (using exponential-plateau and linear fit) for the assessment of individual drug self-administration time curve profiles irrespective of biological sex. We analyzed our data using regression analysis and ANOVA.

**Results:** QSCAn identified three distinct self-administration profiles (consisting of both sexes), which we named exponential-plateau negative (EP-), exponential-plateau positive (EP+), and undefined (EP0). There were no differences in self-administration profiles when we compared males and females within the same group. Differences between sexes (when observed) were due to mismatched comparisons (males from one group versus females from a different group).

**Conclusions:** Our study reinforces the MISSING model for psychostimulant and sucrose self-administration by indicating that differences between males and females (when observed) may not necessarily be driven by biological sex.

**Significance Statement:** Current approaches often interpret variability in psychostimulant self-administration between males and females primarily through the lens of biological sex. However, this framework may overlook meaningful behavioral phenotypes shared across sexes. The present quantitative model suggests that individual patterns of behavior may better account for variability than sex alone, particularly in behaviors not strongly driven by sex-hormone-dependent mechanisms such as drug self-administration. By classifying animals according to behavioral profiles rather than biological sex, this approach may foster the identification of clinically and biologically relevant phenotypes underlying psychostimulant reinforcement.

## Introduction

Psychostimulant use disorders affect both males and females, emphasizing the importance of understanding the role of biological sex in addiction-related behaviors. Consequently, incorporation of sex as a biological variable (SABV) has become a major focus of NIH-funded biomedical research (1–10). While SABV has identified clear sex differences in hormone-related and reproductive behaviors, findings regarding psychostimulant-related behaviors remain inconsistent, with studies reporting both differences and similarities between males and females. Specifically, some studies have reported sex differences in psychostimulant (methamphetamine) self-administration with females consuming more than males (11–14) or males consuming more than females (15–26), while others have found no significant differences (27–43).

These findings indicate that differences between males and females in psychostimulant-related behaviors are less uniformly observed across studies when compared to sex hormone-related behaviors. Moreover, psychostimulant-related behaviors are more directly linked to neurochemical systems such as dopamine than to sex hormones. Thus, biological sex alone may not fully account for the differences, or the lack thereof, in psychostimulant-related behaviors when we compare males and females.

Although biological sex contributes to certain behavioral differences, substantial variability also exists among individuals within each sex, including in psychostimulant-related behaviors. To account for the variability of individual differences while also accounting for biological sex as a variable, we developed an unsupervised clustering model called the MISSING (Mapping Intrinsic Sex Similarities as an Integral quality of Normalized Groups) model (44–48). This model suggests that individual similarities/differences can be organized into group-types that include individuals of both sexes. Rather than assuming that behavioral variation is primarily determined by biological sex, this framework proposes that by accounting for behavioral group identity, most of the apparent sex differences vanish. Consequently, comparisons between males and females may be influenced by relative distributions of behavioral group types latent within each sex. This interpretation differs from traditional analytical approaches which primarily attribute behavioral differences to biological sex and adds a new perspective. Therefore, empirical validation of this model is required. In this study, we tested this hypothesis that observed sex differences between males and females self-administering psychostimulants and/or sucrose may reflect individual or group behavioral variability, rather than biological sex differences alone.

To understand individuals that are similar or different in the way they self-administer drugs, we have to carefully assess self-administration behavior patterns over time. We developed a new Quantitative Structure of Curve Analytical (QSCAn) model for the assessment of behavioral structures (49). We applied this new model to drug self-administration time course curve data to obtain additional behavioral variables beyond those captured by current analytical models, enabling a more comprehensive characterization of individuals (Figure 1). Based on the structure of the self-administration time course curve, individuals are grouped according to shared behavioral patterns. Once individuals are grouped, we can compare males and females within and between those groups. If self-administration behavior differs between males and females within the same behavioral group, these differences are likely associated with biological sex. However, differences observed between males from one behavioral group and females from another behavioral group, referred to as mismatched comparisons, are more likely due to differences in behavioral patterns rather than biological sex.

**Figure 1:**
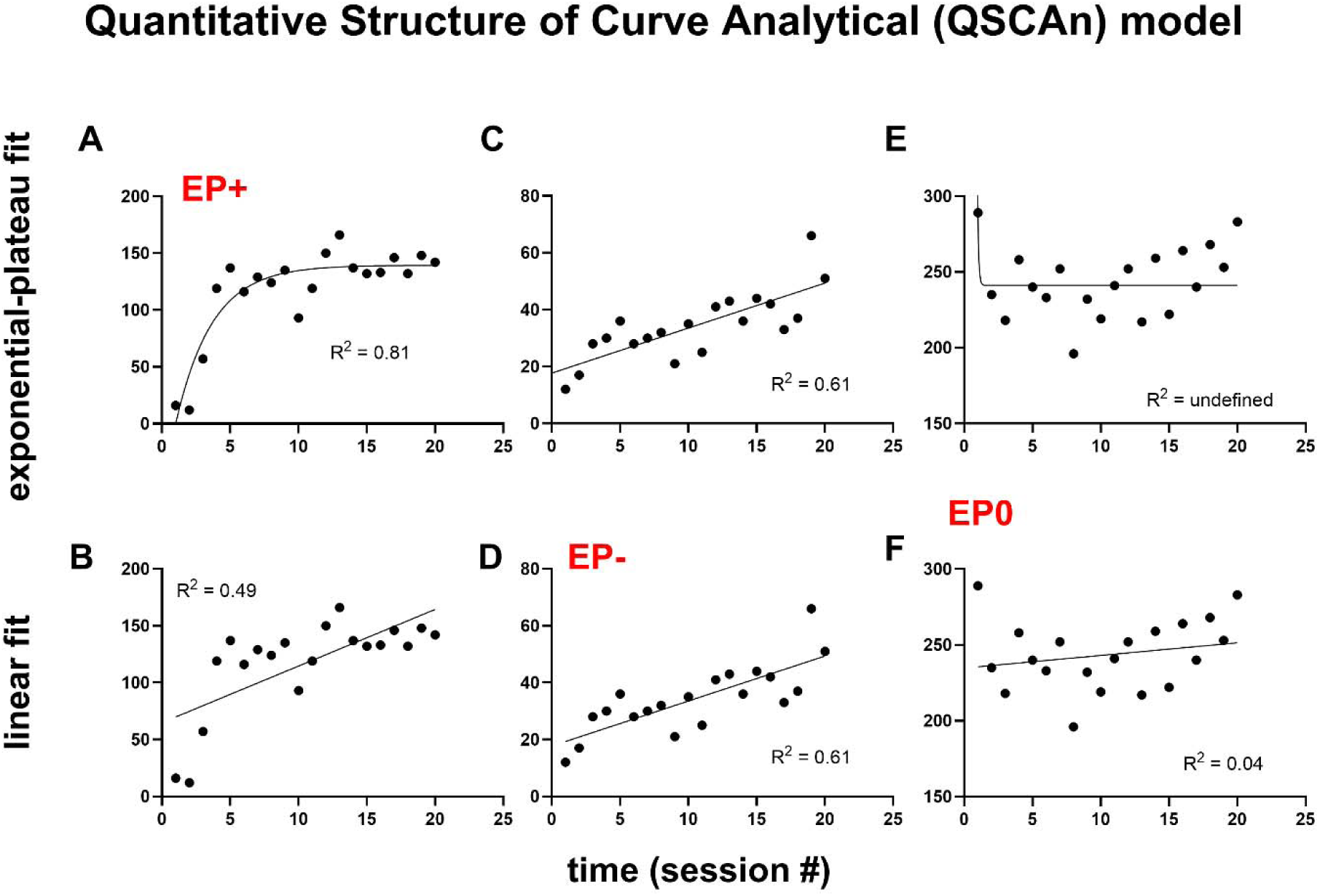
The Quantitative Structure of Curve Analytical (QSCAn) model. The above represent the plots of the self-administration pattern of three individuals: (A-B), (C-D) and (E-F). A and B represent analysis of the self-administration pattern of the same individual with curves being fitted using exponential-plateau (EP) and linear fit, respectively. C and D represent analysis of the self-administration pattern of the same individual with curves being fitted using EP and linear fit, respectively. E and F represent analysis of the self-administration pattern of the same individual with curves being fitted using EP and linear fit, respectively. For individual A-B, the EP fit yielded an R^2^ value for goodness of fit of 0.81 while the linear fit yielded an R^2^ of 0.49. The EP is the better fit for the curve, and this individual (A-B) is termed EP+. For individual C-D, both the EP and linear fit yielded an R^2^ of 0.61, bit the BIC for EP and linear fit were 148.4 and 145.7, respectively. Therefore, the fit with the lower BIC (in this case the linear fit) is the better fit for the curve, and this individual (C-D) is termed EP-. For individual E-F, the R^2^ values for goodness of fit for EP and linear fit were undefined and 0.04, respectively. This curve could neither be fit by EP or linear function and this individual (C-D) is termed EP0.

To test our hypothesis, we allowed different cohorts of male and female Long-Evans rats to self-administer methamphetamine (METH), sucrose, and saline for 20 sessions. Our methods, results, and discussion are described below.

## Methods and Materials

### Animals

All procedures and treatments were approved by the Rowan University Animal Care and Use Committee and by the National Institute on Drug Abuse Animal Care and Use Committee. All procedures and treatments followed the guidelines outlined in the National Institutes of Health (NIH) *Guide for the Care and Use of Laboratory Animals*. The male and female Long-Evans rats were obtained from Charles River Laboratories (CRL, Wilmington, MA) and from the National Institute on Drug Abuse (NIDA). The rats were adults, male and female, age-matched (56-62 days old when received in the housing facility). The rats were housed two per cage on a 12-hour light/dark cycle with free access to food and water.

### Animal use

Overall, 112 rats were tested. There were three major self-administration experiments: saline (n = 3 males, 10 females), METH (n = 25 males, 32 females), and sucrose (n = 20 males, 22 females).

### Experiments

For the METH (and saline) studies, we inserted a catheter into the external jugular vein, as previously described (20). These rats were single-housed after the surgeries to preserve the catheters. All animals were allowed to recover from surgery for approximately 1-2 week(s) before self-administration procedures were initiated. After rats had recovered,, they were allowed to self-administer METH (0.1 mg/kg/infusion or saline) on the FR1 schedule for 6 hours per session for 5 days a week (excluding weekends) for 4 weeks (20 sessions), as described in (20). To minimize the likelihood of infection and the formation of clots, the catheter was flushed daily with 0.2 mL of a sterile solution containing heparin (30.0 IU/mL) and enrofloxacin (5 mg/kg).

For the sucrose studies, no surgical procedures were done. These rats were doubly housed throughout the duration of the experiments. The rats were allowed to self-administer sucrose (one-20 mg pellet, Dustless Precision Pellets, Bio-Serv, Flemington, NJ) on the FR1 schedule for 6 hours per session for 5 days a week (excluding weekends) for 4 weeks (20 sessions).

### The new Quantitative Structure of Curve Analytical (QSCAn) model for the assessment of self-Administration time course

Examples of drug self-administration time course(s) for individuals are shown in Figure 1. This is a plot of time (session #) on the x-axis and corresponding drug consumption on the y-axis. The typical drug self-administration time curve is thought to include an acquisition phase, a plateau phase, and a transition between the acquisition and plateau phases (an inflexion), see Figure 1A. For the typical self-administration time course that includes acquisition and plateau phases, the QSCAn model can track this structure using the exponential-plateau equation to reveal variables that define it. Thus, we employed non-linear regression analysis using the exponential-plateau function (see equation 1) to fit the curve:

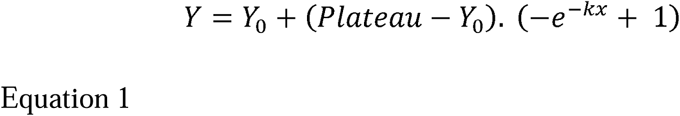

Where plateau is the maximum intake and K is the acquisition rate.

But not all self-administration curves follow an exponential-plateau structure; some have neither an acquisition nor a plateau phase but have an escalation (or linear slope). If an individual self-administration profile can be best fitted using the exponential-plateau function, the individual is grouped as exponential-plateau positive (EP+), see an example in Figure 1A. For the individual in Figure 1A-B, the exponential-plateau function (Figure 1A) revealed a better fit than the linear function (Figure 1B) with respect to goodness of fit R^2^ values. If the better fit is obtained from using the linear function, the individual is termed exponential-plateau negative (EP-).

However, with regards to goodness of fit R^2^ values, some curves can be both EP+ and EP-. To rectify this problem, we obtained the self-administration time curve of all individuals and fit all of them using the QSCAn-exponential-plateau and QSCAn-linear. We obtained the Bayesian Information Criteria (BIC) for these fits and obtained the R^2^ for goodness of fit. The curve with the lowest BIC was selected as the best fit for the self-administration profile. When BIC values were similar, the model with the higher R^2^ was considered the better fit, see an example for the same individual in Figure 1C versus 1D.

In addition, there are subjects that can neither be fit using the exponential-plateau function nor the linear function – these are therefore grouped as undefined (EP0), see an example in Figure 1E-F. Examples of individuals belonging to EP-, EP+, and EP0 are shown in Figure 1.

### Body weight index

We obtained body weights daily throughout the duration of the experiments. There is enough evidence that chronic METH exposure alters feeding behaviors, metabolism, and body weight gain, often producing an initial reduction in bodyweight followed by a slow recovery over time (50–53).

To estimate the effects, if any, of METH and sucrose on body weights, we normalized daily body weights to the initial body weight (body weight on the first day of the experiments) using the formula below:

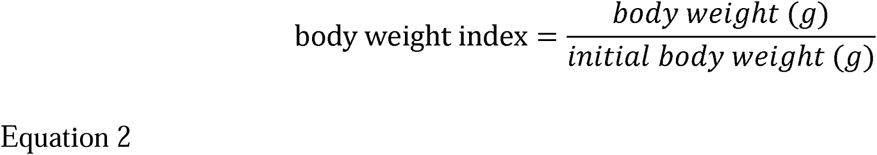

### Statistical analysis

GraphPad Prism v 10 (GraphPad Software, San Diego, CA), SigmaPlot 14.5 (Systat Software Inc., San Jose, CA), and JMP Pro v 18 (SAS Institute Inc., Cary, NC) were employed for statistical analysis. Data were reported as means ± SEM. We conducted QSCAn (exponential-plateau) and QSCAn (linear) for every individual’s self-administration time curve to identify EP+, EP-, and EP0 individuals.

We conducted matched comparisons, comparing males and females within each group. For each study, we used a Two-way repeated-measures ANOVA with sex as a between-subjects factor and time as a within-subjects factor. A sex × time interaction within a group would indicate biological sex differences. We also conducted linear regression analysis to determine if the slope of the relationship between consumption and time was different between sexes. Differences between males and females from the same group are more likely to reflect biological sex differences.

We also conducted mismatched comparisons, which involve comparing males from one group with females from a different group. Self-administration differences between males and females from different groups are likely not due to biological sex but rather due to group-type differences that arise from individual behavior patterns.

Statistical significance was set at P < 0.05 for all analyses, with Tukey’s post hoc test employed when significance was detected.

## Results

### Individual self-administration time curves

As mentioned in the Methods, all graphs were fitted with the QSCAn exponential plateau and the QSCAn linear. Thus, each individual had two graphs, one for QSCAn-exponential plateau and one for QSCAn-linear. The curve with the higher R^2^ and/or the lower BIC was selected as the best fitting model for the time course data. The subjects that were best fitted by QSCAn exponential-plateau were termed EP+. The subjects that were best fitted by QSCAn-linear were termed EP-. Subjects whose self-administration curves could not be fit either by QSCAn-exponential plateau or QSCAn-linear model were classified as EP0 (see Methods). No saline self-administering subjects were EP- or EP+. The composition by sex of each group for METH and sucrose self-administration is shown in Table 1. Note that all groups included individuals from both sexes.

**Table 1:** Composition of self-administration groups by sex METH.

|  | males | females | total |
| --- | --- | --- | --- |

| group |  |  |  |
| --- | --- | --- | --- |
| EP- | 6 | 8 | 14 |
| EP+ | 9 | 21 | 30 |
| EP0 | 10 | 3 | 13 |
| total | 25 | 32 | 57 |

Sucrose
|  | males | females | total |
| --- | --- | --- | --- |
| group |  |  |  |
| EP- | 2 | 2 | 4 |
| EP+ | 15 | 13 | 28 |
| EP0 | 3 | 7 | 10 |
| total | 20 | 22 | 42 |

### Self-administration time curve group comparisons

After grouping individuals as EP-, EP+, and/or EP0 (and including saline group), we conducted a mixed-effects model (Two-way) repeated measures ANOVA (factors are group and time). With dependent variables as drug infusions, Two-way repeated measures ANOVA revealed a group x time interaction (F 20.06, 435.7 = 10.70, P < 0.0001, Figure 2A) – Tukey’s post hoc tests revealed that saline and EP0 were not different at any time point. Also, EP+ and EP-were not different at any time point. However, saline was different from EP+ at all time points except time point 1, whereas saline was different from EP-at all time points except time points 1-3.

**Figure 2:**
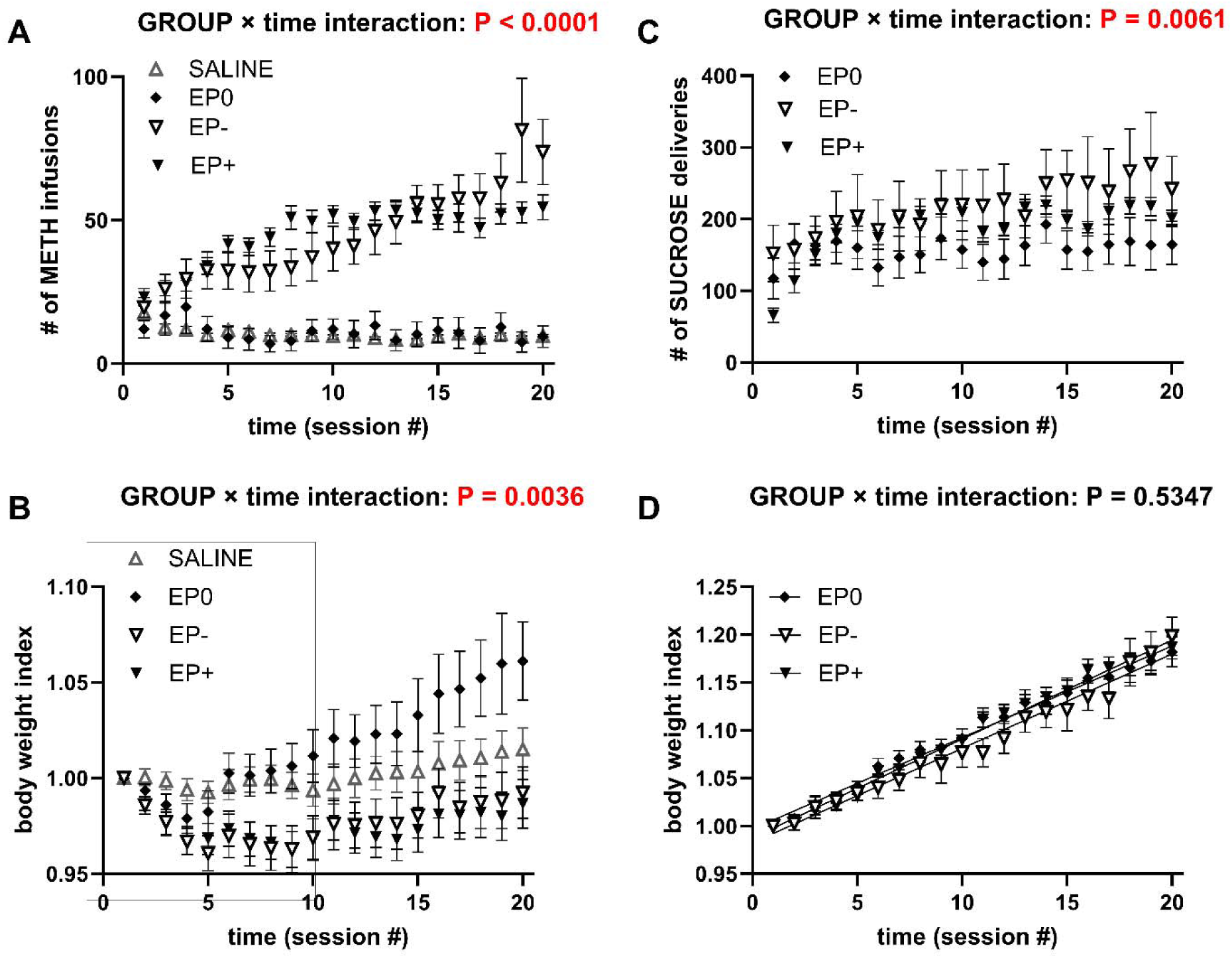
Identification of distinct behavioral groups composed of both sexes using QSCAn analysis of METH and sucrose self-administration. The individuals were classified into EP+, EP−, and EP0 groups, each containing both males and females. (A) METH self-administration across sessions. (B) Body weight index across METH self-administration sessions. (C) Sucrose self-administration across sessions. (D) Body weight index across sucrose self-administration sessions. Two-way repeated-measures ANOVA detected significant group × time interactions for METH intake (A), METH body weight index (B), and sucrose intake (C), but not sucrose body weight index (D). Data are mean ± SEM.

When we compared the time curves of only EP- and EP+, we detected a group x time interaction (F 5.931, 246.0 = 7.660, P < 0.0001) – Importantly, the distinction between EP- and EP+ was also detected by conventional repeated-measures ANOVA, indicating that the differences identified by QSCAn are reflected in the raw self-administration data.

With body weight index as the dependent variable, Two-way repeated measures ANOVA revealed a group x time interaction (F 6.146, 135.2 = 3.376, P = 0.0036, Figure 2B) – Tukey’s post hoc tests revealed that saline and EP0 were not different at any time point. Also, EP+ and EP-were not different at any time point. However, saline was different from EP+ at all time points from 1 – 9, except for time point 6. Saline was different from EP-at time points 4 and 5 only. EP-and EP+ were not different. Within group comparisons showed that EP-group body weight index was different from time point 1 only for time point 4 whereas for EP+ body weight index, there was a decrease at time points 2, 4, 5 and 9 relative to time point 1. Because the body weight index time course appeared to be a biphasic curve, we compared the slopes (regression analysis) of body weight index over time for days 1-10 and days 11-20 (Figure 2B). For the slope of the plot of body weight index over time for days 1-10, linear regression analysis revealed a significant difference between at least one group from the others (F 3, 692 = 6.098, P = 0.0004). Further analysis revealed the following: EP-v EP+ (P = 0.9710), EP-v EP0 (P = 0.0004), EP-v saline (P = 0.0328), EP+ v EP0 (P = 0.0005), EP+ v saline (P = 0.0598) and EP0 v saline (P = 0.0363). For slope of the plot of body weight index over time for days 11-20, there were no differences between groups (F 3, 692 = 1.151, P = 0.3278).

For the sucrose time curve (Figure 2C), we obtained the following results: group × time interaction (F 8.347, 162.8 = 2.762, P = 0.0061). Linear regression revealed the following slopes (significant difference from zero) for EP- (5.436 ± 1.679, F 1, 78 = 10.49, P = 0.0018), EP+ (4.490 ± 0.4875, F 1, 558 = 84.84, P < 0.0001), and EP0 (0.9981 ± 1.061, F 1, 198 = 0.8857, P = 0.3478). Slope comparisons revealed significant differences between at least one of the groups (F 2, 834 = 6.264, P = 0.0020). Further analysis for slope revealed the following: EP-v EP+ (P = 0.5104), EP-v EP0 (P = 0.0261), EP+ v EP0 (P = 0.0008). A comparison between linear slopes of EP-v EP+ revealed no differences between these groups (P > 0.05), however when we conducted nonlinear regression with exponential fit, we detected significant differences. Non-linear regression revealed the following: plateau (F 1, 634 = 13.46, P = 0.0003) and K (F 1, 634 = 6.351, P = 0.0120). Importantly, the distinction between EP- and EP+ was not detected by conventional repeated-measures ANOVA and linear regression but was detected by nonlinear regression (exponential-plateau) that followed the QSCAn structure. This indicates that the differences identified by QSCAn are reflected in the raw self-administration data.

With body weight index as the dependent variable, Two-way repeated measures ANOVA did not reveal any group x time interaction (F 4.376, 85.34 = 0.8061, P = 0.5342, Figure 2D). The plot of body weight index over time was linear. For slope of the plot of body weight index over time for days 1-20, regression analysis revealed no significant difference between groups (F 2, 834 = 0.9229, P = 0.3978, Figure 2D).

### Comparison of sexes without accounting for distinct self-administration time-curve groups: CURRENT model

For METH consumption levels (Figure 3A), mixed-effects repeated measures ANOVA (with SEX and time as factors) revealed no SEX × time interaction (F 4.599, 249.8 = 0.6703, P = 0.6338). Linear regression analysis revealed no male versus female differences in the linear relationship between consumption and time (F 1, 1123 = 1.315, P = 0.2517). Non-linear regression analysis also revealed that there are no differences between males and females for K (F 1, 1121 = 0.3498, P = 0.5543) and plateau (F 1, 1121 = 0.04418, P = 0.8336).

**Figure 3:**
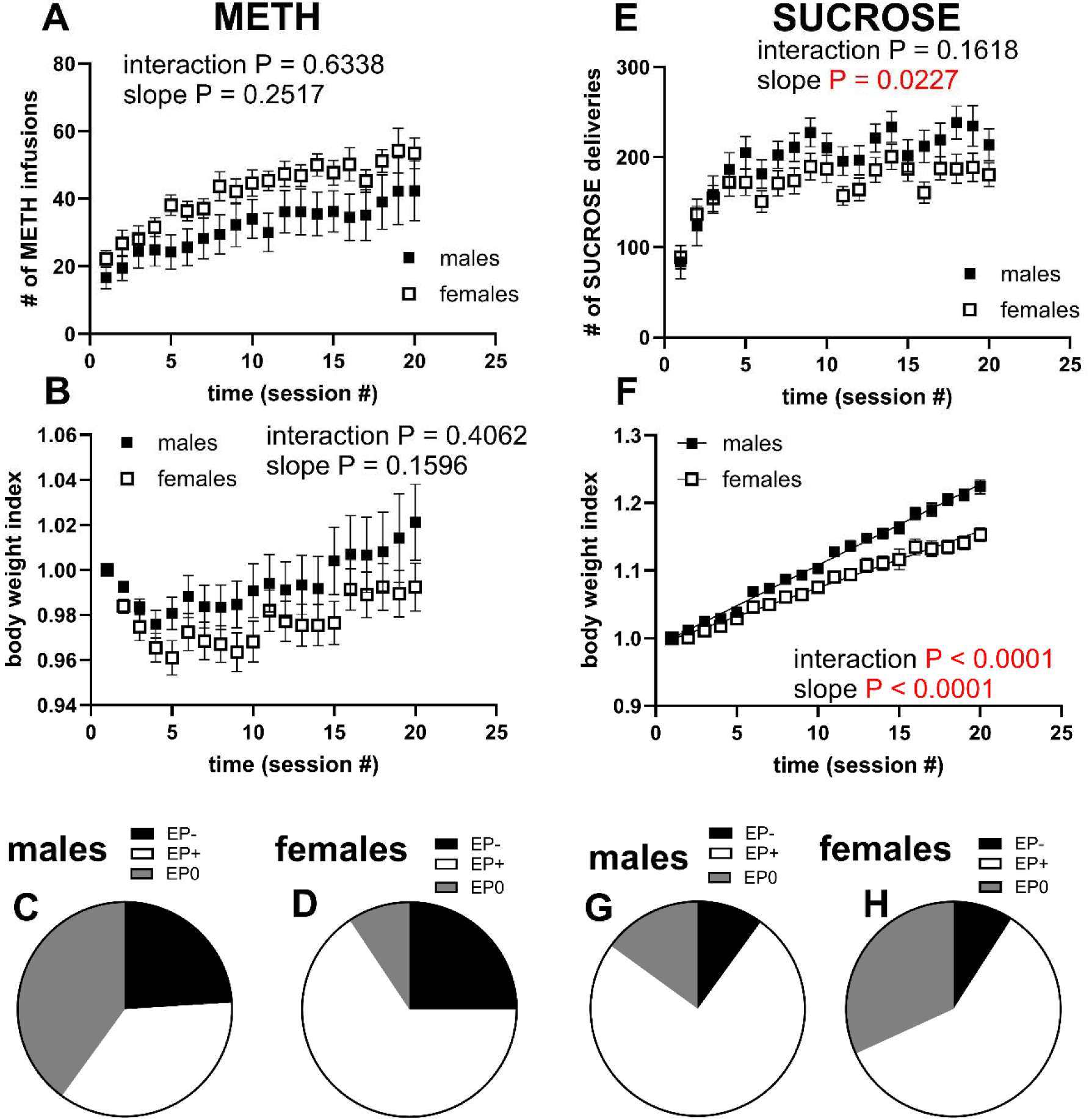
There were no sex differences for METH self-administration but there were sex differences for sucrose self-administration (when we do not account for the distinct groups of males and females within the data set). Graph A-B represent a plot of consumption over time and body weight index over time, respectively, for METH, with each plot showing males (closed squares) versus females. Graph C-D represent the relative composition of each of the identified behavioral groups (EP-, EP+, EP0) for METH self-administration with respect to males and females. Graph E-F represent a plot of consumption over time and body weight index over time, respectively, for sucrose, with each plot showing males (closed squares) versus females. Graph G-H represent the relative composition of each of the identified behavioral groups (EP-, EP+, EP0) for sucrose self-administration with respect to males and females. There was no SEX by time interaction (or differences in linear slope) for METH consumption (A) and body weight index changes due to METH consumption (B). There was no SEX by time interaction, but there were sex differences in differences in linear slope for sucrose consumption time course (E). There was a significant SEX by time interaction and a significant difference in linear slope for male versus female sucrose body weight index-time course (F).

For the effect of METH on body weight index (Figure 3B), mixed-effects repeated measures ANOVA revealed no SEX × time interaction (F 1.806, 99.31 = 0.8859, P = 0.4062). Linear regression analysis revealed no male versus female differences in the relationship between body weight index and time (P = 0.2517). Note that the comparison between males and females (Figure 3A-B) was a comparison between 3 distinct groups that included males and females (Fig 3C-D), see group composition by sex in Table 1.

For sucrose (Figure 3E), mixed-effects repeated measures ANOVA revealed no SEX × time interaction (F 4.175, 167 = 1.648, P = 0.1618). Linear slope comparisons (male slope = 4.816 ± 0.7099 versus female slope = 2.779 ± 0.5549) revealed a significant difference (F 1, 836 = 5.208, P = 0.0227). Non-linear regression revealed male versus female differences for plateau (F 1, 834 = 11.81, P = 0.0006), but not for K (F 1, 834 = 0.6023, P = 0.84379).

For the effect of sucrose on body weight index (Figure 3F), mixed-effects repeated measures ANOVA revealed a significant SEX × time interaction (F 2.688, 107.5 = 10.62, P < 0.0001). Linear regression analysis revealed a significant difference between sexes in the slope of the relationship between body weight index and time (F 1, 836 = 72.58, P < 0.0001). Note that the comparison between males and females (Figure 3E-F) was a comparison between 3 distinct groups that each included males and females (Fig 3G-H), see group composition by sex in Table 1.

### Comparison of the drug intake time course of males and females while accounting for the distinct self-administration time-curve groups: the MISSING model

The statistics for the METH and sucrose groups for drug intake time course are shown in Tables 2 and 3, respectively. For METH, there were no differences when we conducted comparisons between males and females within the same drug self-administration time-curve behavioral profiles (Figure 4A, E and I). When there were differences, it was because we conducted mismatched comparisons, see Figure 4C-D, F-H. For sucrose, there were no differences when we conducted comparisons between males and females within the same self-administration time-curve behavioral profiles (Figure 5A, E and I). As with METH, when there were differences between sexes, it was because we conducted mismatched comparisons, see Figure 5G-H.

**Table 2:**
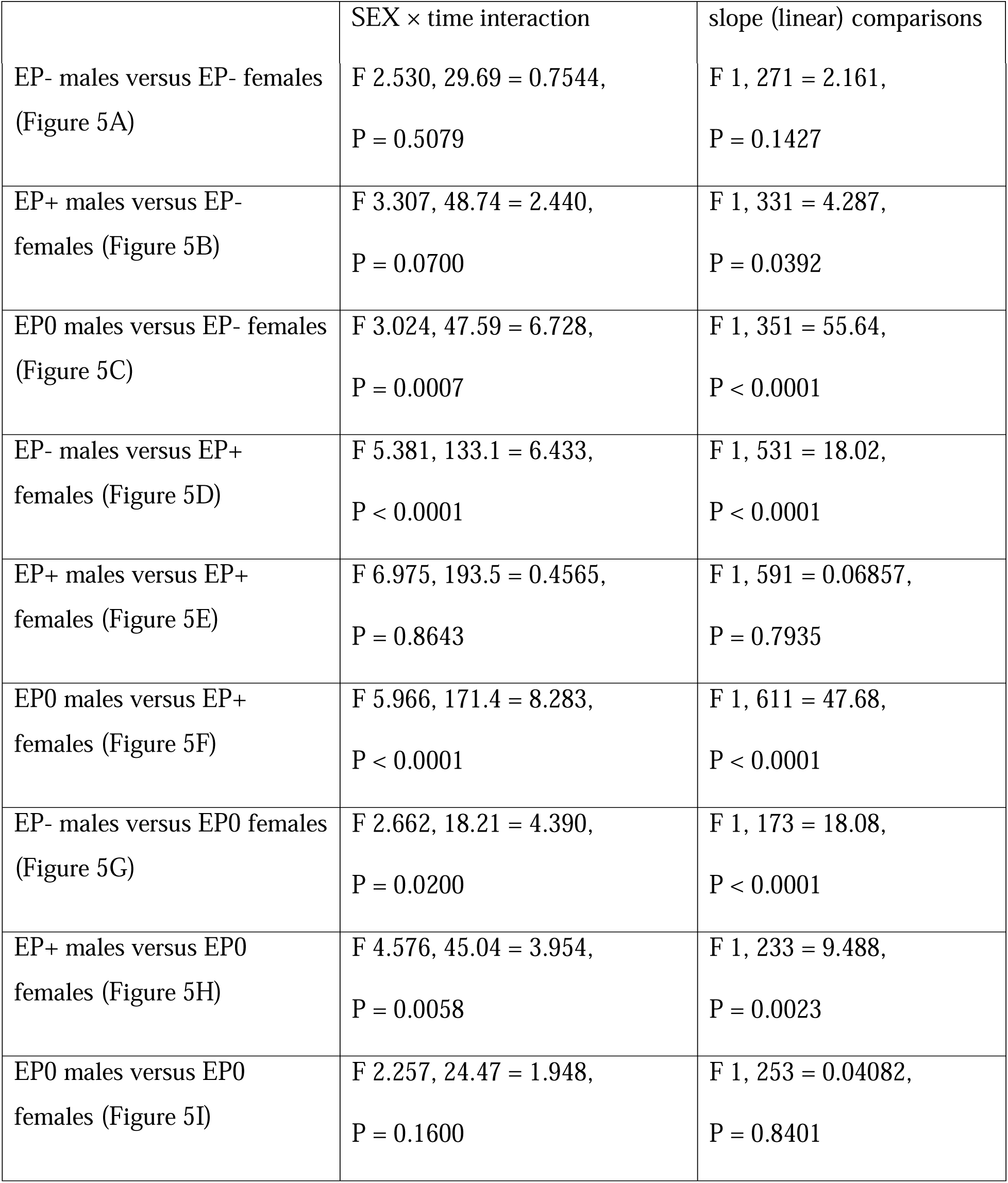
Statistics for the MISSING model for METH groups (males versus females): drug intake time course.

| | SEX × time interaction | slope (linear) comparisons |
| --- | --- | --- |
| EP- males versus EP- females<br>(Figure 5A) | F 2.530, 29.69 = 0.7544,<br>P = 0.5079 | F 1, 271 = 2.161,<br>P = 0.1427 |
| EP+ males versus EP-<br>females (Figure 5B) | F 3.307, 48.74 = 2.440,<br>P = 0.0700 | F 1, 331 = 4.287,<br>P = 0.0392 |
| EP0 males versus EP- females<br>(Figure 5C) | F 3.024, 47.59 = 6.728,<br>P = 0.0007 | F 1, 351 = 55.64,<br>P < 0.0001 |
| EP- males versus EP+<br>females (Figure 5D) | F 5.381, 133.1 = 6.433,<br>P < 0.0001 | F 1, 531 = 18.02,<br>P < 0.0001 |
| EP+ males versus EP+<br>females (Figure 5E) | F 6.975, 193.5 = 0.4565,<br>P = 0.8643 | F 1, 591 = 0.06857,<br>P = 0.7935 |
| EP0 males versus EP+<br>females (Figure 5F) | F 5.966, 171.4 = 8.283,<br>P < 0.0001 | F 1, 611 = 47.68,<br>P < 0.0001 |
| EP- males versus EP0 females<br>(Figure 5G) | F 2.662, 18.21 = 4.390,<br>P = 0.0200 | F 1, 173 = 18.08,<br>P < 0.0001 |
| EP+ males versus EP0<br>females (Figure 5H) | F 4.576, 45.04 = 3.954,<br>P = 0.0058 | F 1, 233 = 9.488,<br>P = 0.0023 |
| EP0 males versus EP0<br>females (Figure 5I) | F 2.257, 24.47 = 1.948,<br>P = 0.1600 | F 1, 253 = 0.04082,<br>P = 0.8401 |

**Table 3:** Statistics for the MISSING model for Sucrose groups (males versus females): drug intake time course.

| | SEX × time interaction | slope (linear) comparisons |
| --- | --- | --- |
| EP- males versus EP- females<br>(Figure 6A) | F 1.679, 3.358 = 0.6403,<br>P = 0.5563 | F 1, 76 = 0.5215,<br>P = 0.4724 |
| EP+ males versus EP- females (Figure 6B) | F 3.326, 49.89 = 0.8285,<br>P = 0.4949 | F 1, 336 = 0.1886,<br>P = 0.6343 |
| EP0 males versus EP- females<br>(Figure 6C) | F 1.137, 3.412 = 0.1351,<br>P = 0.7653 | F 1, 96 = 0.9741,<br>P = 0.3262 |
| EP- males versus EP+ females (Figure 6D) | F 4.335, 56.36 = 1.817,<br>P = 0.1334 | F 1, 296 = 2.061,<br>P = 0.1522 |
| EP+ males versus EP+ females (Figure 6E) | F 3.546, 99.29 = 1.766,<br>P = 0.1492 | F 1, 556 = 3.035,<br>P = 0.0821 |
| EP0 males versus EP+ females (Figure 6F) | F 2.216, 31.03 = 1.089,<br>P = 0.3543 | F 1, 316 = 1.403,<br>P = 0.2371 |
| EP- males versus EP0 females<br>(Figure 6G) | F 3.595, 25.17 = 3.153,<br>P = 0.0352 | F 1, 176 = 4.944,<br>P = 0.0275 |
| EP+ males versus EP0 females (Figure 6H) | F 4.097, 81.94 = 4.394,<br>P = 0.0027 | F 1, 436 = 12.42,<br>P = 0.0005 |
| EP0 males versus EP0 females (Figure 6I) | F 1.567, 12.54 = 0.3032,<br>P = 0.6918 | F 1, 196 = 0.08196,<br>P = 0.7750 |

**Figure 4.**
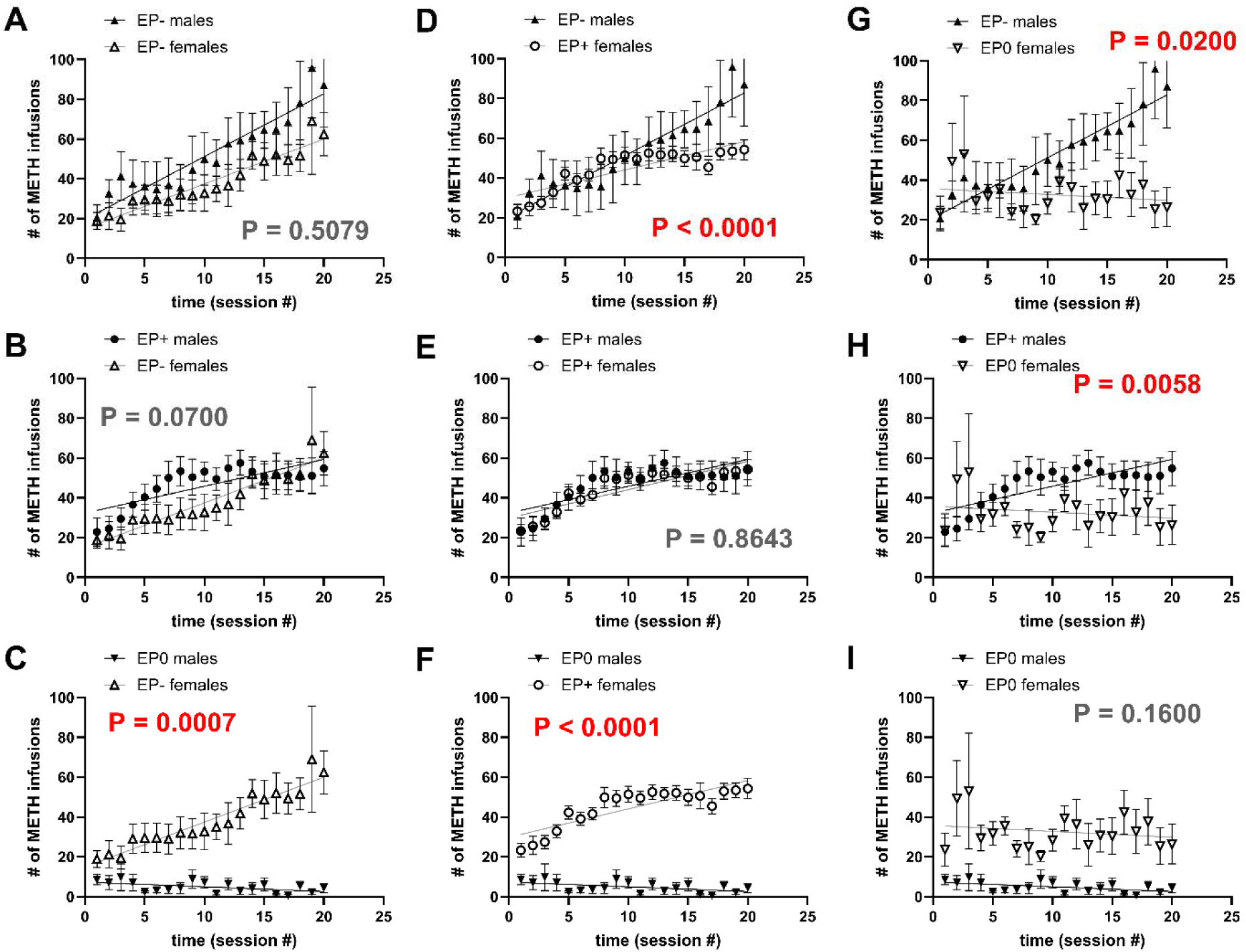
MISSING model for METH self-administration time curve groups: drug intake time course. Graphs A and E show matched comparisons between males and females belonging to the same behavioral group (EP-, EP+, EP0, respectively) B-D, and F-H show mismatched comparisons between males and females belonging to different behavioral groups. P values represent the sex x time interaction from two-way repeated-measures ANOVA.

**Figure 5.**
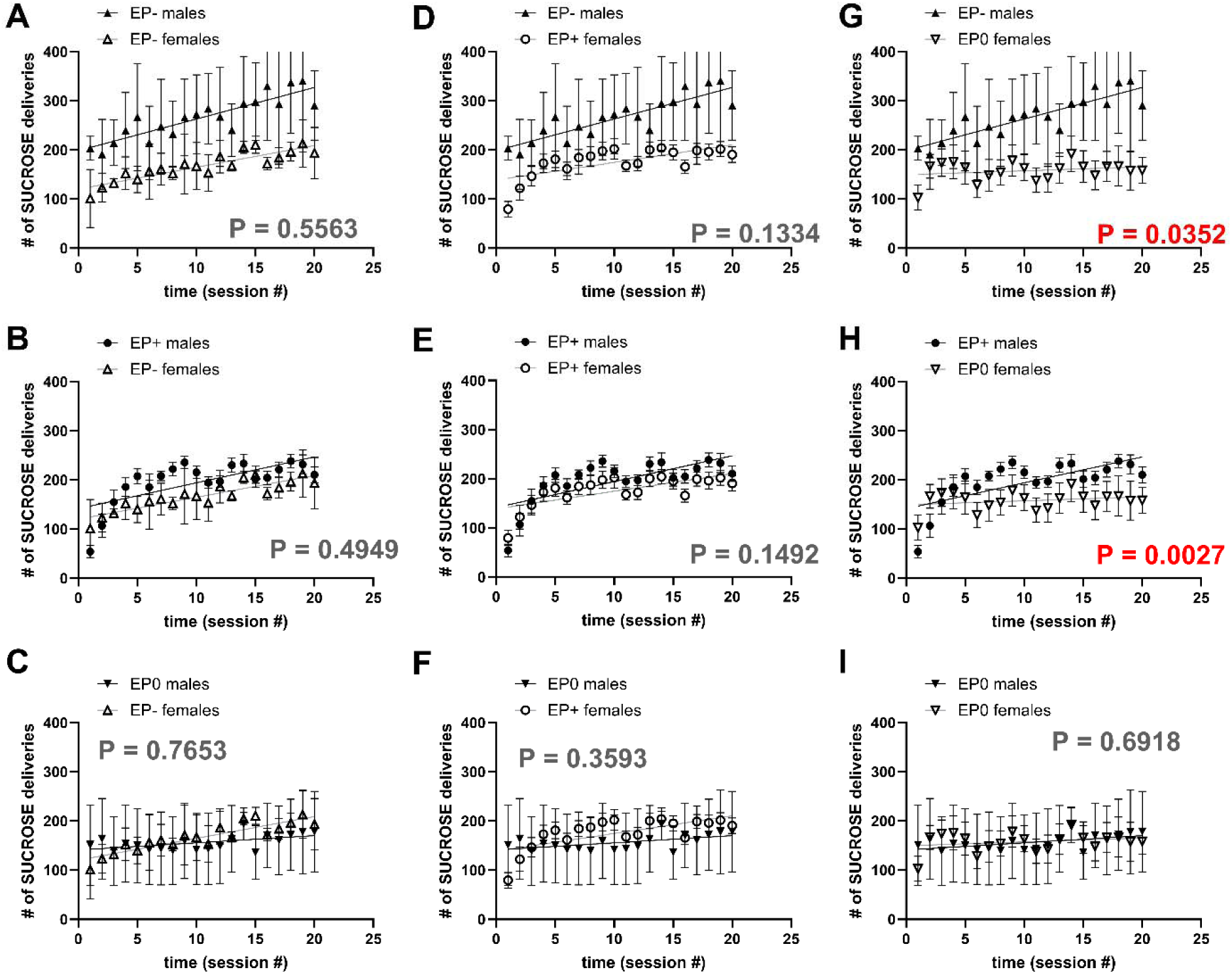
MISSING model for sucrose self-administration time curve groups: drug intake time course. Graphs A, E, and I show matched comparisons between males and females belonging to the same behavioral group (EP-, EP+, EP0, respectively). B-D, and F-H show mismatched comparisons between males and females belonging to different behavioral groups. P values represent the sex x time interaction from two-way repeated-measures ANOVA

### Comparison of the body weight index time course of males and females while accounting for the distinct self-administration time-curve groups: the MISSING model

The statistics for the METH and sucrose groups for body weight index time course are shown in Tables 4 and 5, respectively. For METH, there were no differences when we conducted comparisons between males and females within the same drug self-administration time-curve behavioral profiles for body weight index time course (Figure 6A, E and I). When there was a difference, it was because we conducted mismatched comparisons, see Figure 6F.

**Table 4:**
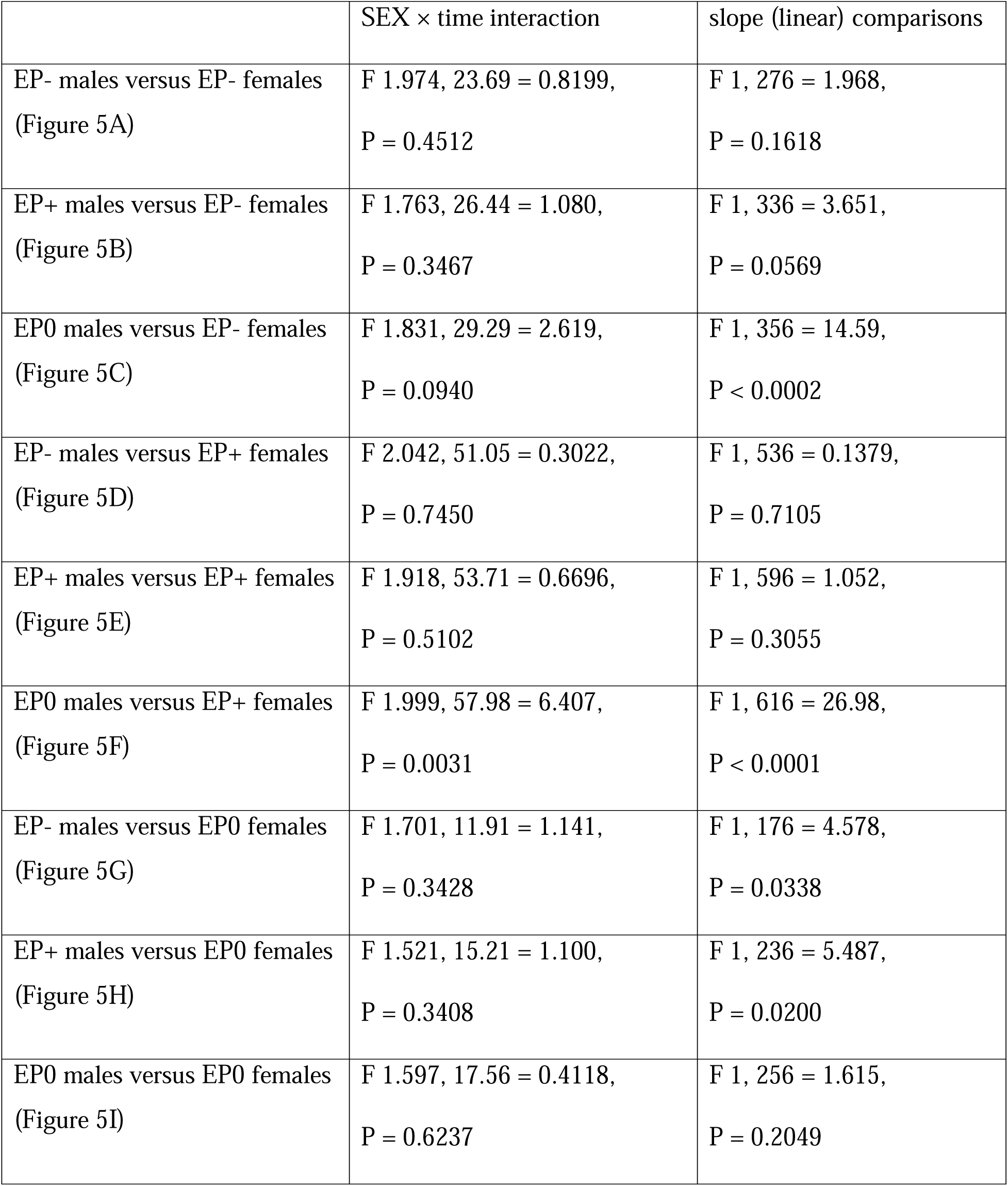
Statistics for the MISSING model for METH groups (males versus females): body weight index time course.

**Table 5:** Statistics for the MISSING model for Sucrose groups (males versus females): body weight index time course.

| | SEX × time interaction | slope (linear) comparisons |
| --- | --- | --- |
| EP- males versus EP- females<br>(Figure 6A) | F 1.627, 3.253 = 2.273,<br>P = 0.2365 | F 1, 76 = 11.99,<br>P = 0.0009 |
| EP+ males versus EP- females (Figure 6B) | F 1.582, 23.73 = 3.973,<br>P = 0.0410 | F 1, 336 = 17.19,<br>P < 0.0001 |
| EP0 males versus EP- females<br>(Figure 6C) | F 1.829, 5.487 = 2.394,<br>P = 0.1793 | F 1, 96 = 8.892,<br>P = 0.0036 |
| EP- males versus EP+ females (Figure 6D) | F 2.703, 35.14 = 1.361,<br>P = 0.2711 | F 1, 296 = 7.334,<br>P = 0.0072 |
| EP+ males versus EP+ females (Figure 6E) | F 2.602, 67.65 = 8.649,<br>P = 0.0001 | F 1, 556 = 62.38,<br>P < 0.0001 |
| EP0 males versus EP+ females (Figure 6F) | F 2.722, 38.11 = 1.142,<br>P = 0.3416 | F 1, 316 = 8.394,<br>P = 0.0040 |
| EP- males versus EP0 females<br>(Figure 6G) | F 1.626, 11.38 = 1.598,<br>P = 0.2425 | F 1, 176 = 4.551,<br>P = 0.0343 |
| EP+ males versus EP0 females (Figure 6H) | F 1.634, 32.69 = 5.642,<br>P = 0.0115 | F 1, 436 = 31.11,<br>P < 0.0001 |
| EP0 males versus EP0 females (Figure 6I) | F 1.618, 12.95 = 0.9609,<br>P = 0.3899 | F 1, 196 = 4.418,<br>P = 0.0368 |

**Figure 6.**
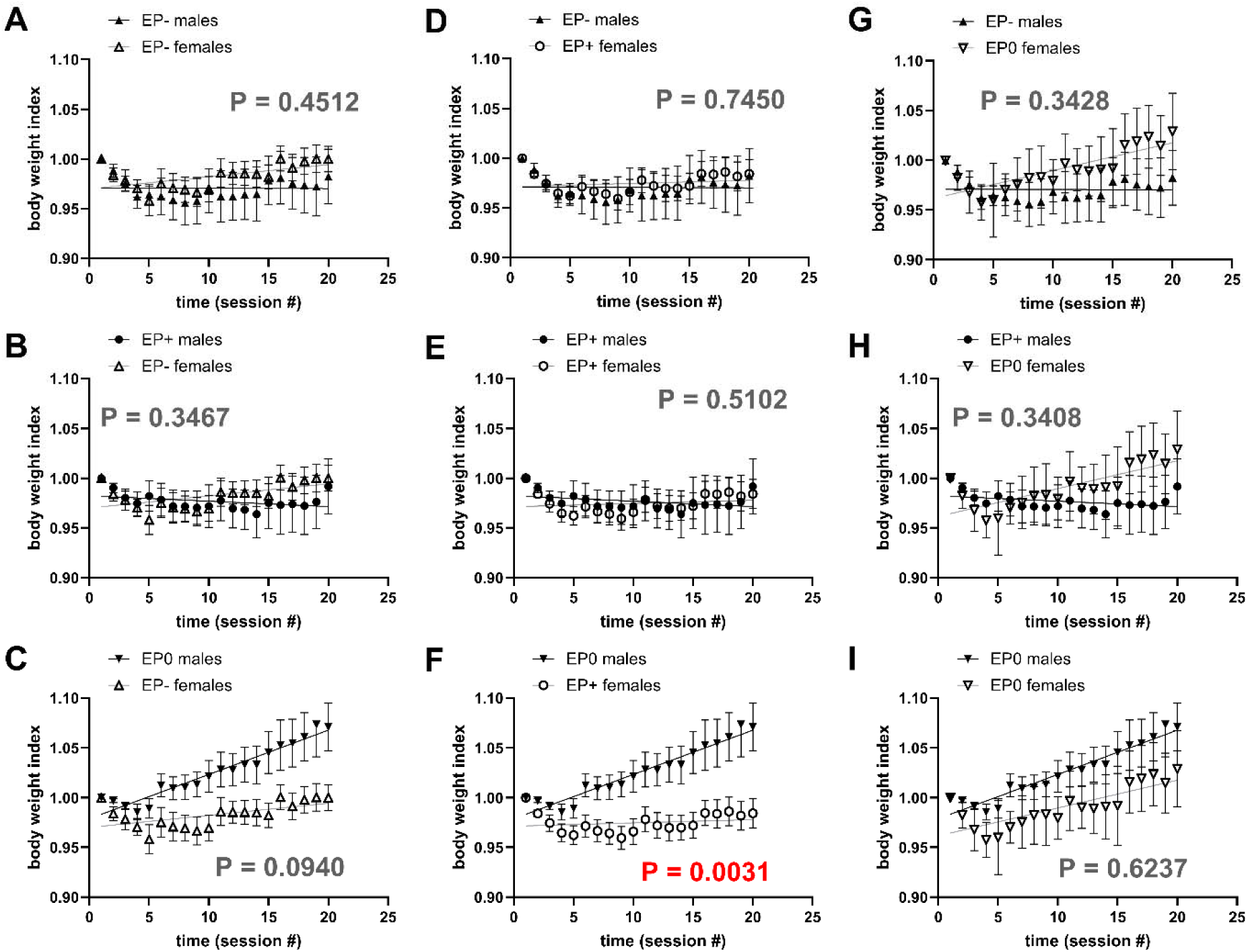
MISSING model for METH self-administration time curve groups: body weight index time course. The graphs are comparisons of 1) males versus females belonging to the same group (matched comparisons for graphs A, E and I for EP-, EP+ and EP0, respectively), and 2) males versus females belonging to different groups (mismatched comparisons for EP+ males versus EP-females, EP0 males versus EP-females, EP-males versus EP+ females, EP0 males versus EP+ females, EP-males versus EP0 females, and EP+ males versus EP0 females in graphs B, C, D, F, G, and H, respectively). Note that there were no differences for matched comparisons. When there was a significant difference between sexes, it was between males and females from distinct groups (F). These differences are not biological sex-related differences – they are group-related differences. The P values written into the graphs are for SEX by time interaction.

Interestingly, for body weight index time course for sucrose, we detected significant differences between males and females in the EP+ group (Figure 7E). We also detected differences between males and females following mismatched comparisons: EP+ males v EP-females (Figure 7B) and EP+ males v EP0 females (Figure 7H).

**Figure 7.**
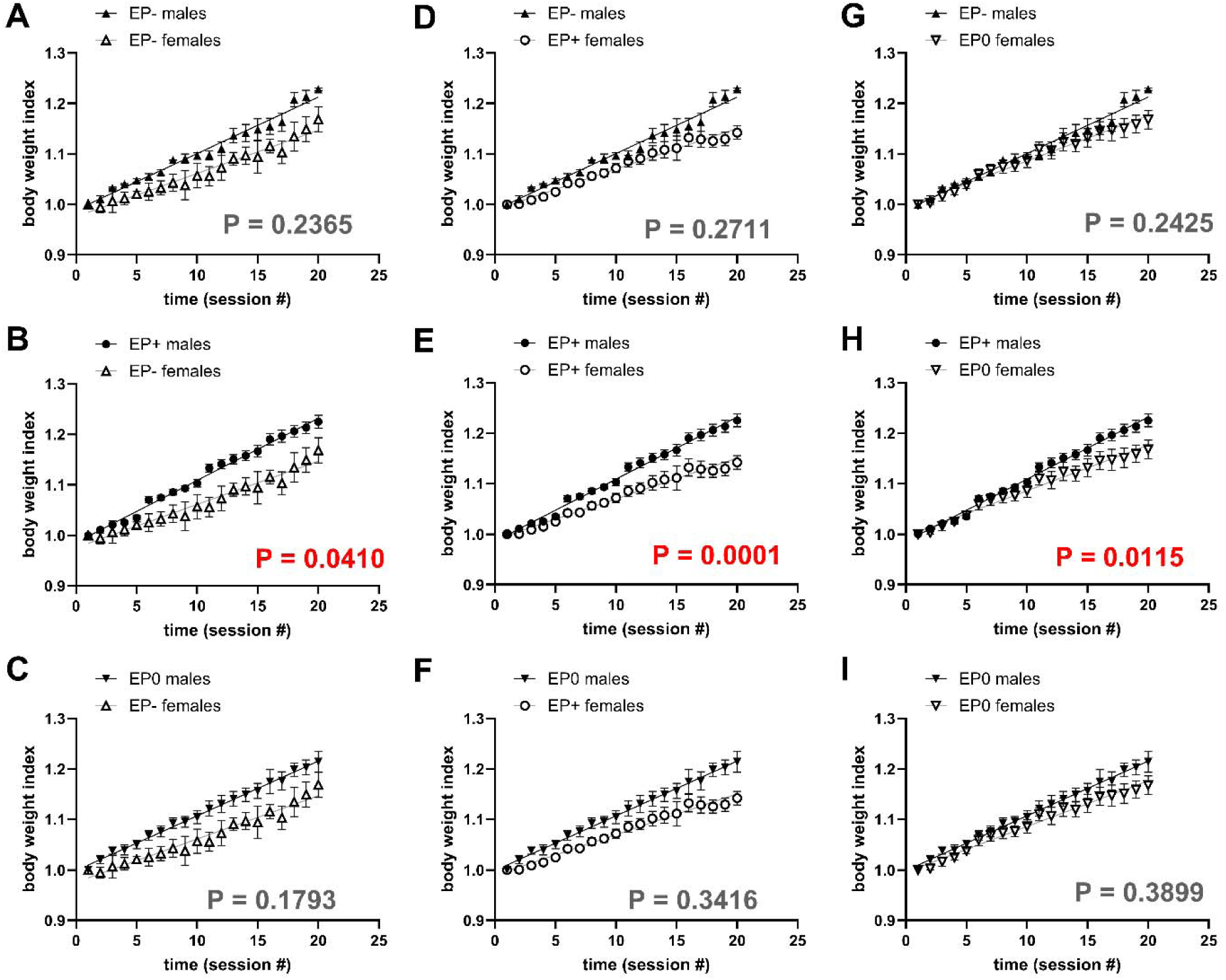
MISSING model for sucrose self-administration time curve groups: body weight index time course. The graphs are comparisons of 1) males versus females belonging to the same group (matched comparisons for graphs A, E and I for EP-, EP+ and EP0, respectively), and 2) males versus females belonging to different groups (mismatched comparisons for EP+ males versus EP-females, EP0 males versus EP-females, EP-males versus EP+ females, EP0 males versus EP+ females, EP-males versus EP0 females, and EP+ males versus EP0 females in graphs B, C, D, F, G, and H, respectively). Note that there was a significant difference between males and females in EP+ group (E). There were also differences for some mismatched comparisons (see B and H). The differences between males and females from the matched comparison represent a biological sex difference. The differences between males and females from the mismatched comparison do not represent a biological sex difference. The P values written into the graphs are for SEX by time interaction.

## Discussion

The primary goal of this study was to evaluate the MISSING model for psychostimulant self-administration, as we have previously done for psychostimulant-induced locomotor activity (45,46,48). We employed a new QSCAn model for the assessment of individual self-administration time curves (Figure 1) just as we have done for psychostimulant self-administration dose-response curves (49). From QSCAn of METH and sucrose self-administration time curves, we identified three groups containing both sexes, which we named EP-, EP+, and EP0 (Table 1).

The QSCAn model appears more sensitive than conventional analytical approaches in detecting differences in self-administration trajectories. This was particularly evident for sucrose self-administration, where repeated measures ANOVA and linear regression failed to distinguish EP-and EP+ groups, whereas the QSCAn model identified significant differences in both K and plateau. These findings suggest that reliance on conventional analyses which primarily evaluate differences in average responding across time or between groups, may obscure meaningful differences in behavioral trajectories. By modeling the shape of self-administration curves, QSCAn can detect variation in behavioral patterns that is not always captured by ANOVA or linear regression.

An additional implication of these findings is that vulnerability to psychostimulant self-administration and reinforcement may not be uniform across individuals. Although all animals had same self-administration conditions, QSCAn identified a sub-group (EP0) that persistently showed low levels of responding, little to no escalation of intake, and behavioral trajectories that closely resembled saline controls than the other METH-taking groups. However, EP0 animals were not indistinguishable from saline animals. Bodyweight analysis revealed differences between EP0 and saline groups despite their similar intake patterns. Note also that the composition of this group (EP0) relative to all subjects was not insignificant – they made up 22.8% and 23.8% of METH and sucrose consumers, respectively, see Table 1. While the biological significance of this finding remains unclear, it suggests that EP0 animals may still be affected by METH in ways that are not captured by self-administration measures alone. These findings suggest that access to a psychostimulant doesn’t necessarily mean equal reinforcement across all individuals (54,55) and that substantial heterogeneity may exist in the degree to which METH can function as a salient reinforcer.

Additionally, EP0 group may potentially be used as a comparison group for future studies examining the neurobiological consequences of psychostimulant self-administration. Unlike saline and sucrose controls, EP0 animals were exposed to the same drug, drug-associated cues and catheterization as EP- and EP+ animals yet exhibited different intake trajectories. Traditional control groups like saline, sucrose, and yoked controls each address important questions but differ from active METH-taking animals in reinforcement contingencies, drug exposure, and route of administration. EP0 subjects may provide a unique opportunity to distinguish biological changes associated with active METH-taking behavior from those associated with experimental exposure conditions alone. However, additional studies are necessary to determine the utility of this group as a control and to establish whether it represents a distinct biological phenotype.

Importantly, the EP0 group included both males and females (like EP- and EP+ groups), suggesting that this variability is not sex-specific. Consistent with MISSING model, these findings highlight the importance of examining individual behavioral patterns before grouping them by sex, as population averages may obscure meaningful differences in vulnerability to drug-taking behavior.

An important utility of MISSING model is in its ability to distinguish apparent sex differences that arise from behavioral heterogeneity from sex differences that persist after behavioral group identity is accounted for. In this study, in most of the cases, significant differences between males and females appeared when making mismatched comparisons, suggesting that behavioral group identity contributed substantially to the observed sex effects. However, for sucrose body weight index, EP+ males and EP+ females showed a significant difference from each other suggesting that MISSING model can not only identify apparent sex differences driven by behavioral heterogeneity but can also reveal sex differences that persist within the same behavioral group. This sex difference could therefore be more likely driven by biological sex.

Taken together, these findings have three major implications for the preclinical psychostimulant self-administration research. First, they support the MISSING model by showing that meaningful behavioral groups can emerge independent of biological sex and that apparent sex differences arise from comparisons between different behavioral groups rather than sex itself. Second, they highlight the importance of trajectory-based approaches such as QSCAn which can identify differences in behavioral dynamics obscured by group averages and/or linear trends. Finally, the identification of EP0 group suggests that vulnerability to psychostimulant reinforcement may not be uniform across individuals and that some subject may never acquire drug self-administration despite equivalent access conditions to the drug. Together, these findings emphasize the importance of examining individual behavioral trajectories before assigning biological interpretations and suggest that incorporating these behavioral trajectories into analyses may improve our understanding of individual variability and sex differences in psychostimulant reinforcement.

There are few limitations to the study that must be considered. First, the present findings are based on behavioral analysis, and future studies including neurobiological measures are necessary to determine the biological basis of identified groups. Second, housing conditions were different among experimental groups. Sucrose animals were group-housed while METH and saline animals were single-housed, which may have influenced the behavioral outcomes.

In conclusion, our study supports the MISSING model as a psychostimulant self-administration paradigm and explains the inconsistencies in the literature regarding the observation of sex differences in psychostimulant self-administration. The MISSING model reveals that when we observe differences between males and females in psychostimulant self-administration, such differences are likely not due to biological sex but to individual/group behavioral patterns. Our conclusions represent a paradigm shift from the *status quo* and may significantly advance the field of sex differences research.

## Acknowledgements

The authors acknowledge the support of NIH grants **DA054461** (MOJ), **DA055701** (TMK), and **GM136492** (AJ, TMK). The content is solely the responsibility of the authors and does not necessarily represent the official views of the National Institutes of Health. Additional support was provided by the Intramural Research Program of the National Institute on Drug Abuse (NIDA IRP), Rowan University via the Camden Health Research Initiative grant #1050 (TMK), the Francis R. Lax Fund for Faculty Development (MOJ), and faculty startup funds (MOJ).

